# Phenotypic plasticity and evolution of thermal tolerance in two lineages of bacteria from temperate and hot environments

**DOI:** 10.1101/2020.08.22.262865

**Authors:** Enrique Hurtado-Bautista, Laura F. Pérez-Sánchez, África Islas-Robles, Gustavo Santoyo, Gabriela Olmedo-Álvarez

## Abstract

Despite the crucial role of microorganisms to sustain life on Earth, there is little research on the evolution of thermal tolerance of bacteria in the face of the challenge that global warming poses. Phenotypic adaptation to a new environment requires plasticity to allow individuals to respond to selective forces, followed by adaptive evolution. We do not know to what extent phenotypic plasticity allows thermal tolerance evolution in bacteria at the border of their physiological limits. We analyzed growth and thermal reaction norms to temperature of strains of two bacterial lineages, *Bacillus cereus sensu lato* and *Bacillus subtilis sensu lato*, that evolved in two contrasting environments, a temperate lagoon (T) and a hot spring (H). Our results showed that despite co-occurrence of members of both lineages in the two contrasting environments, norms of reactions to temperature exhibited a similar pattern only within the lineages, suggesting fixed phenotypic plasticity. Additionally, within the *B. cereus* lineage, strains from the H environment showed only two to three °C more heat tolerance than strains from the T environment. The limited evolutionary changes towards an increase in heat tolerance in bacteria should alert us of the negative impact that climate change can have on all biological cycles in the planet.

## Introduction

Temperature is one of the most important physical factors that define a species fundamental niche (1). It affects many phenotypes, and numerous investigations on adaptation have focused on temperature to understand how it impacts physiological processes at the molecular level (2,3). Temperature affects a broad range of phenotypes, so it is used as a model to investigate how phenotypic plasticity evolves. Understanding phenotypic plasticity has become of high importance, given the expected temperature rise in the planet. Studies in ectotherm groups have suggested that variation in upper thermal limits is narrower compared to that of lower temperature and have suggested that evolution of heat tolerance is constrained. This asymmetry has been reviewed for terrestrial endo- and ectotherms, insects, amphibians and plants (4,5), and more recently an extensive data set was analyzed by (6).

In contrast to the many studies that have been done in eukaryotes to determine their thermal plasticity, in bacteria, there are few examples. Unlike the restrictions to the temperatures where eukaryotic organisms can thrive, Archaea and Bacteria can be found in extreme environments, from freezing (−40 °C) (7), to very high (50 and to 100 °C) (8). Their ubiquitous occurrence does not mean, however, that individual phylum or species have a broad spectrum of tolerance to temperature. Like eukaryotic ectotherms, individual bacterial taxa exhibit a limited temperature niche.

Phenotypic plasticity is the ability of an organism to exhibit distinct phenotypes when exposed to different environments (9,10), and allows organisms to acclimate to changes, extending the ecological range of a species, so they can survive exposure to pressures and creating the opportunity for assimilation (Waddington 1953, cited by (11). Assimilation is the mechanism through which the initial plastic response allows diversification through genetic changes that stabilize the expression of the induced phenotype (12). Interaction among plasticity, life history and evolution persist for generations (13).

Organisms’ genetics is the basis of phenotypic plasticity and the degree to which an organism can alter its phenotype, partly governed by functional genomic mechanisms, will contribute to delimiting the range of environmental conditions to which it can acclimate. If an individual’s biological response to a changing environment is a function of gene content and its regulation, it could be expected that genetically close organisms that experience similar environmental pressure may exhibit similar plasticity to respond to that particular stress. However, when species encounter changes in their environment, long term persistence will require the evolution of their plasticity. Since some habitat will, in fact, be less favourable to fitness, costs and limits to the evolution of phenotypic plasticity are expected (14).

Temperature is a chosen variable in many studies that evaluate patterns of growth rate, survival, reproduction and doubling time in the population of bacteria. Growth rate represents a simple response variable of continuous phenotypes (15). Phenotypic plasticity can be evaluated through reaction norms (10,16, and Wolterek, 1909, cited by (11)). Reaction norms are a description on how a phenotype varies as a continuous function of the environmental cues and is represented by a curve on a graph that plots a phenotype against an environmental factor (Figure 1a). In a historical account on the study of norms of reaction, (11) cites Dobzhansky’s writing: “what changes in evolution is the norm of reaction of the organism to the environment”. The complete reaction norm is a trait, and thus may be different between genotypes; it is genetically variable and thus, it can evolve.

**Figure 1.**
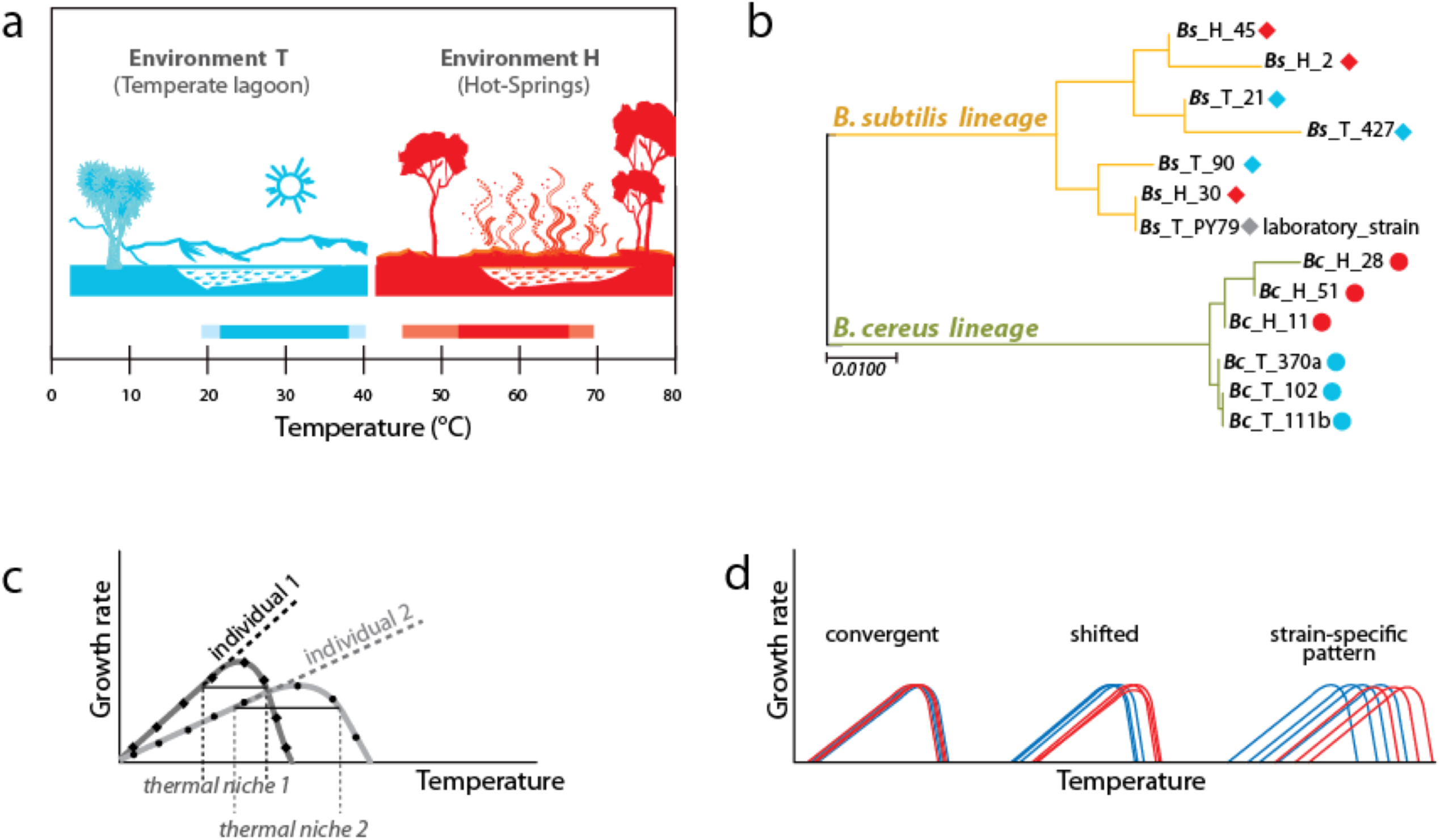
Experimental approach to analyzing phenotypic plasticity in two lineages of bacteria from contrasting environments. a. Environments of isolation. Laguna Intermedia of Churince, Cuatrocienegas (temperature range of 17 to 40 °C) and Hot-springs in Michoacán (Mexico) (temperature range between 48 and 70 °C). b. Phylogenetic relationships for the Bacillus lineages representing mesophilic (M) and thermotolerant (T) life histories. B. subtilis and B. cereus from a Temperate-lagoon (blue), Hot-spring (red) and a laboratory strain (PY79) (extense phylogeny in supplementary Fig. 2). c. A norm of reaction represents performance of individuals under different environmental conditions. The amplitude of the curve is the breath of the performance and thermal niche is the rank between the two x-values of 75% of maximal growth at optimal temperature (3). d. Scenarios of phenotypic plasticity: individuals from different lineages or environments may exhibit the same response (convergence), a shifted response of the isolates from the hot-spring, or even strain-specific patterns regardless of genetic lineage (blue lines, environment T; red lines, environment H).

Only 0.3% of the studies on plasticity-led evolution have been done in bacteria (17). The few studies on phenotypic plasticity to temperature in bacteria have been carried out on model laboratory bacteria, such as *Bacillus subtilis* (18) and *Escherichia coli* (see, for example, 19–21). However, these strains may not be optimal to capture the complexity of plasticity, as many traits may have been lost through passages under laboratory conditions. Genetic analyses have revealed genetic variation for thermotolerance under laboratory conditions (22 and references therein), but thermal plasticity of bacteria as a result of selection pressures in nature remains largely unknown.

The *Bacillus* genus is characterized by endospore-forming bacteria, and representatives of this genus are present in almost every wild environment around the world (23–25). *Bacillus* is an interesting model to study phenotypic plasticity. Its ability to develop a highly resistant spore allows survival at a temperature that would be lethal for the vegetative cell, thus allowing it to survive extreme changes. How then could refinement of its phenotype to tolerate higher temperature occur if the immediate response of these bacteria to stress was sporulation? The fact that the *Bacillus* can tolerate heat in their sporulated form does not make the *Bacillus* thermophiles, as most species cannot grow above 50°C. Some *Bacillus* species have been recovered from extreme environments and are thermophiles, such as *Bacillus infernus* and *Bacillus fumarolis* (26), but the best-studied thermophilic genus in the Firmicutes are usually classified in different genera, such as *Geobacillus, Thermaerobacter*, and *Thermobacillus* (27).

It has been of interest to understand the capability of a given species to occupy different thermal niches. Hot springs have been recurrent systems for investigating niche diversification in natural communities of microorganisms, and thermophilic bacteria thrive in these environments. Weltzer and Miller showed that *Chloroflexus* strains from the White creek thermal gradient have diverged in the temperature range for growth (28). On the other hand, laboratory strains of the *Synechococcus* A/B group of cyanobacteria isolated from different temperatures from both Yellowstone and Oregon hot springs are ecological specialists with divergent temperature ranges for growth (29).

Although some *Bacillus* strains representatives of mesophilic clades are sometimes isolated from hot-springs, there is typically little information of their taxonomy and even of their temperature tolerance. It is possible that, with a few exceptions, many strains in the *Bacillus* genus isolated from hot-springs are not thermophilic and they tolerate heat as spores. For instance, seldom are mesophilic *Bacillus* species, such as *Bacillus cereus* and *Bacillus subtilis*, recovered from hot springs. Being so ubiquitous, can they extend their range of temperature tolerance and evolve into thermophilic strains?

Among the numerous *Bacillus* species recognized, some lineages have been extensively studied, such as the *B. cereus sensu lato*, that includes *Bacillus cereus, Bacillus thuringiensis*, and *Bacillus anthracis* ((30), and the *Bacillus subtilis* complex, that includes *Bacillus subtilis, Bacillus amyloliquefaciens, Bacillus licheniformis*, and *Bacillus pumilus*, among others (27,31). The most recent study by (24) through the use of 700 conserved genes, showed that the *B. cereus* and *B. subtilis* lineages form two distinct clades. At genome level, genome size and gene content are distinct in these two lineages. *B. cereus* typically possesses a genome of between 5 to 6 mega base pairs, while the genome of *B. subtilis sensu lato* is around 4 mega bases long (32).

Bacteria, in general, exhibit considerable genetic variability, in part from their ability to interchange genes through horizontal gene transfer. Within the genus *Bacillus* there is a large intraspecies phenotypic variability (33) and a significant variation in the genetic repertoire through microevolution (34). Up to 30% of genes may be different within bacterial species (35). Organisms that have evolved in a given environment may be constrained in their response, maybe from having adjusted their genes to their particular environment (24). If this was the case, their life history could come very close to constitute its genetic history as well. Bacteria are excellent models to explore the evolution of plasticity, through the evaluation of reaction norms to temperature. Their genetic variability makes them special cases to explore whether the genetic mould is so malleable that their norms of reaction change to adjust to the environment or if, on the contrary, despite this variability, their reaction norm is fixed, such that it can be a trait of the phylogeny. At the molecular level, temperature response has been extensively studied in bacteria and particularly the response elicited by both cold- and heat shock (36,37). We do not know, however, whether the large repertoire of genes required for thermal adaptation constrains the evolution of tolerance.

In this work, we evaluated phenotypic plasticity to thermal tolerance in a lineage Vs. environment model in bacteria from natural settings. By examining the evolution of upper thermal limits in bacterial strains from contrasting environments, it is possible to evaluate trait limits related to evolutionary history. The bacterial strains used in this study comprised two lineages within a genus, *B. cereus sensu lato* and *B. subtilis sensu lato*. The strains were obtained from a hot-spring (environment H) and a temperate lagoon (environment T), both in Mexico, and were used to address the following questions: Do individuals of closely related lineages with a similar history of temperature selection (either in the hot-spring or in the temperate lagoon) exhibit convergence in their norms of reaction? Do the Bacillus from the hot-springs evolve tolerance to temperature in their vegetative stage?

Our results showed that reaction norms to temperature of the different individuals reflected their evolutionary history. The *B. cereus* and *B. subtilis* lineages each exhibited distinct response patterns, suggesting that the genetic architecture of each lineage constrained their phenotypic plasticity despite their sharing of environmental conditions. For both lineages, covariation was observed between environmental temperature and thermal tolerance phenotype, suggesting temperature adaptation. The individuals from the hot-springs were, as expected, more tolerant to hot temperature, yet, their tolerance did not match the hot-springs temperature suggesting, particularly for the *B. cereus* lineage, that its ecological strategy depends mainly on sporulation. These results may suggest that sporulation decreases the opportunity for evolving tolerance and that the lineage in its vegetative state is already close to its thermal tolerance limit.

## Materials and Methods

### Evaluation of mesophile and thermophile strains

Bacteria classified as mesophilic can tolerate a range of 18 to 45 °C, while thermotolerant bacteria tolerate from 22 to 60 °C. Both mesophilic and thermotolerant bacteria have growth optima below 50 °C. Thermophilic bacteria, in contrast, have an optimal growth temperature above 60 °C (38). Mesophilic *Bacillus* were collected from the Churince water system, where daily and seasonal variation in temperature have been recorded, since the spring is fed by subterranean water and the system is held in a range of 31°C near the water spring and closer to ambient air temperature, 18-31 °C, in the faraway limits of the lagoon system (39). The thermotolerant Bacillus strains in this study were collected in the geothermal system of the Araro region, located in the central part of Mexico, inside the trans-Mexican volcanic belt located in the Michoacan state. The dominant bacteria in this extreme environment were firmicutes, inhabiting the microbial mats in the springs (40). Temperature and physicochemical parameters were evaluated in different seasons and found to fluctuate between 45 and 55 °C (Bonita spring) and 63 o 74 °C (Tina hot-spring) (40). For this study, we chose sets of strains from two closely related taxa, both of the *Bacillus* genus (as explained below). Six were isolated from the Temperate intermediate lagoon (environment T) and six more from the hot-spring in Michoacan (environment H) (Fig. 1a). We included a *B. subtilis* laboratory strain, PY79, presumably mesophilic (41). Bacterial strains were kept in frozen stocks at −70 °C. To observe the phenotype of their colonies selected strains were streaked out on semisolid Marine medium and incubated for 24 h to 48 h at 37, 44, 50 and 55 °C (Photographs of some of the plates are shown in Supplementary Fig. 1).

### Strain selection from 16S rRNA phylogenetic reconstruction analysis

PCR of 16S rRNA genes was obtained from a collection of strains from the temperate lagoon and the hot-springs. Forward and reverse sequences were obtained by Sanger dideoxy sequencing, edited by cutting off low-quality segments and concatenated as a consensus sequence for each gene using Bioedit version 7.0.5.3. (See Supplementary Fig. 2). Phylogenetic analysis was done in Mega 7.0.26, after alignment using Muscle. We chose six strains that grouped with the *B. cereus sensu lato* (were 99 % similar based on sequence variation of the 16S rRNA gene) and six strains from the *B subtilis* lineage. For the simplified phylogeny shown in Fig. 1b, gene alignment of the 16S rRNA gene of the 13 chosen strains was carried out using Muscle and tree construction was performed by the Maximum Likelihood method with the HKY+G substitution model using MEGA version 7.0.26 (42).

**Figure 2.**
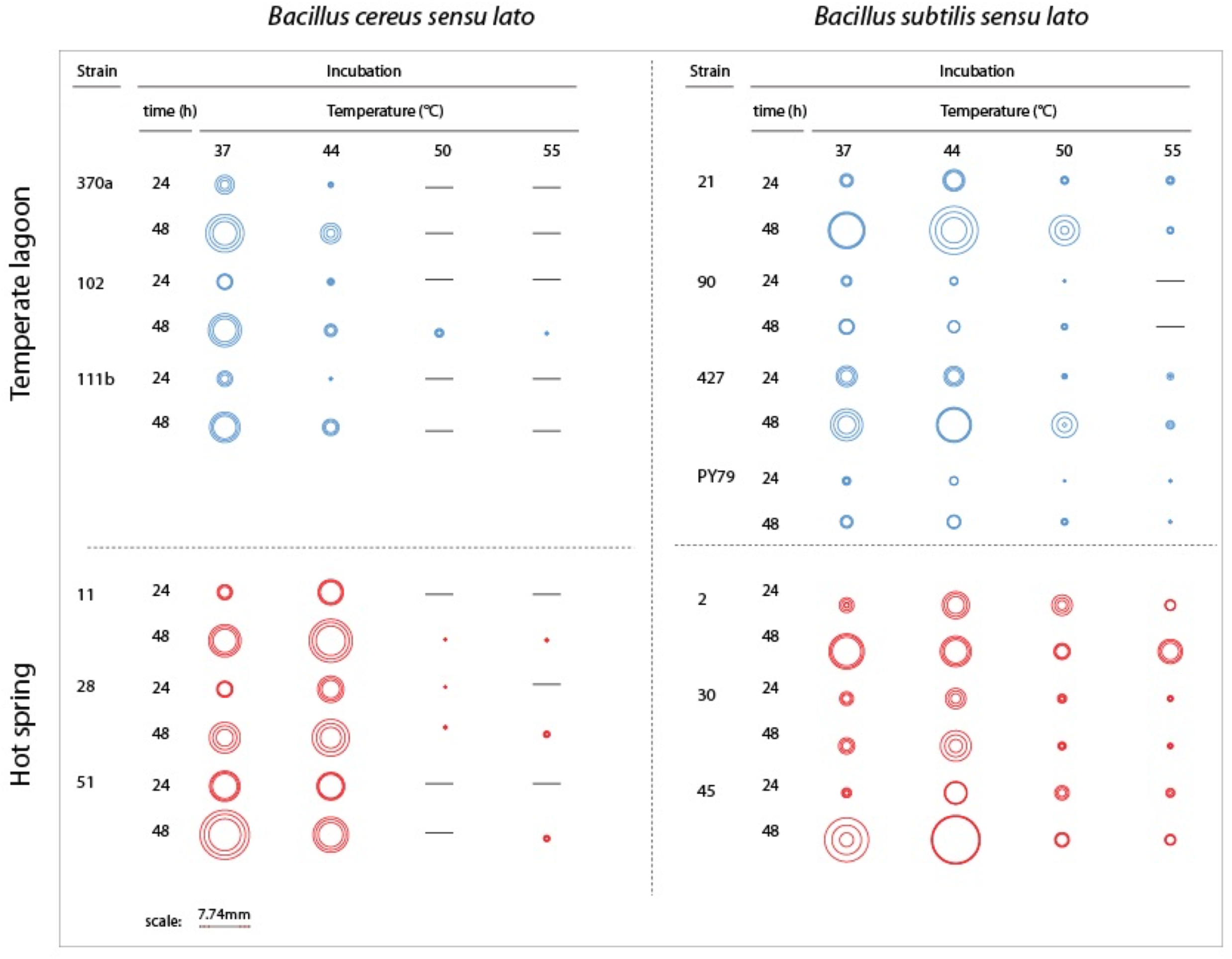
Growth of colonies of strains from the two lineages and environments at different temperatures. Graphical representation of the size of single colonies after 24 and 48 h incubation at different temperatures (37, 44, 50 and 55 °C) on semisolid Marine Medium. Three isolated colonies were chosen, and their size represented in a circle; the inner circles and the outer circles represent, respectively, the lower and upper limits of the standard deviations error bars. Dashes indicate that no growth was observed. Supplementary Fig. 1 shows photographs from some of the plates.

### Determination of growth rates evaluation and norms of reaction

Bacterial isolates from −70°C stocks were reactivated on semisolid Marine Medium (43) at 37°C for 20 hrs. One colony from each strain was inoculated into to 50 ml Falcon tubes with 5 ml of Marine medium-broth and incubated overnight at 37°C in a shaking incubator. After 18 hours of incubation, an aliquot of 50 μl from the cultures, was transferred to tubes with fresh Marine-broth and incubated two more hours to bring them to exponential growth condition, before measurement of the growth kinetics curve. Five μl of each culture was used for inoculating 200-well microtiter plates of the Bioscreen C (Labsystems, Helsinki, Finland) previously filled with 175 μl of fresh Marine medium-broth. Measurements were carried out with a 420-580 nm filter, with three replicas. Optical density was measured every 30 min for 20 h. For reaction norms to temperature we obtained kinetic curves at 17, 27, 37, 41, 43, 46, 49 and 55°C. Doubling time was calculated using an exponential model of growth for building the reaction norms for each temperature and evaluated the difference among groups with a t-Test (supplementary figures). Multiple statistical Anovas (0.05 of significance) for comparison throughout all the entire reaction norms were performed in Statgraphics version 15.2.06. The optimal temperature was defined as that with the maximum peak in growth rate throughout the range of tested temperatures. The thermal niche was calculated as the range of temperatures over which the observed doubling rate equaled or exceeded 75% of the peak doubling rate (3). For the evaluation by groups of reaction norms, we combined a set of results from the reaction norm for every temperature value. While comparing species we grouped data of growth rate at each temperature of *B. subtilis* strains from both environments and compared against the grouped *B. cereus* data. To compare environment we grouped the growth rate data of each temperature of *B. subtilis* plus *B. cereus* from each one of the places of origin. An ANOVA in R package 3.6.2 (with significance level at 0.05) to identify statistical differences in the double comparison.

## Results

### Evaluation of phenotypic plasticity in a *Bacillus* two-lineages model, each with members that evolved in contrasting temperature environments

We studied *Bacillus* isolates in a classical gene X environment setup, using strains isolated from sediment in the Intermediate lagoon of Churince in Cuatrocienegas, Coahuila (43) and isolates cultivated from the mats of hot-springs in Michoacán (40). The two different environments appear to have non-overlapping temperature ranges. The temperate water and sediment of the Churince system, from which part of our microbial collection was obtained, has a temperature that fluctuates between 18 to 36 °C (we refer to this as Environment T), while that of the hot spring fluctuates between 45 and 70 °C (Environment H) (Fig. 1a). *Bacillus* strains are easily recovered from both the T and H environments. The strains used in this work have been previously reported (40,43). Phylogenies based on 16S rRNA gene of several strains from the different environments were obtained to select those that would be genetically closest (Supplementary Fig. 2). Emphasis was made in clades *B. subtilis sensu lato* and *B. cereus sensu lato*. These lineages are referred to as Bc and Bs, for short. We chose three strains from each *Bacillus* lineage and from each environment (H and T) to evaluate phenotypic plasticity through comparative norms of reaction. We also included in the study a laboratory strain of *B. subtilis*, strain PY79 (41). A simplified phylogeny is shown in Fig. 1b.

### Growth and colony size differences between the Bc and Bs lineages challenged at high temperature

Colony growth was evaluated on semisolid marine medium with incubation at different times and temperatures (Fig. 2 and Supplementary Fig. 2). We observed growth and colony size differences between strains from the Bs and Bc lineages. Regardless of the environment of isolation, the Bs lineage strains were more tolerant to high temperature than those of the Bc lineage, although the size of single colonies was generally smaller than those of the Bs lineage. The strains from the Bs lineage from the environment T can still grow at 50 °C, although forming small colonies. In two of these strains Bs-T-427 and the laboratory strain PY79, some growth can be observed even at 55 °C. Strains from the Bc lineage, in contrast, cannot grow at 50 °C. The strains from the Bc lineage from environment H exhibited perceptibly larger colonies than their counterparts from environment T, and even at 44 °C grew robustly, suggesting adaptation of these strains to growth at this temperature. Noteworthy, even when challenged at higher temperature (44 °C), colony growth was sustained, since colony size after 48 h incubation was noticeably larger than at 24 h incubation (see Fig. 2 for a schematic of colony size).

### Distinct norms of reaction to temperature of the two Bacillus lineages that co-occur in the Churince temperate lagoon

Phenotypic plasticity was assayed through the norm of reaction to temperature for each strain. Growth curves obtained temperatures from 17 to 55 °C. The thermal niche of each strain was calculated as the range of temperatures over which the observed doubling rate equaled or exceeded 75% of the peak doubling rate (3) (Fig. 1c). In this gene for environment evaluation different norm of reaction scenarios were possible (Fig. 1c and 1d): In one scenario (fixed plasticity), bacteria from both lineages could exhibit the same response to temperature, regardless of the environment where they had evolved. In a second scenario, a shift of tolerance towards higher temperature in both lineages would be observed. In this last case, the selective environmental pressure would result in a convergent phenotypic response regardless of the lineage. In this scenario, strains from the Bs and Bc lineage would exhibit the same response to temperature within each environment. In a third scenario, even if strains from environment H tolerated higher temperature than those from environment T, each individual would exhibit dissimilar norms of reaction to temperature, regardless of the lineage.

Fig. 3a shows the profiles of the norms of reaction to temperature for the strains from the temperate lagoon (T environment). It is observed that despite sharing the same environment, the norms of reaction of the two lineages did not converge. The individuals from the T environment of the Bc and Bs lineage had norms of reaction with a distinct pattern, clearly different from one another. All Bc strains exhibited a higher growth rate at temperatures from 17 to 40 °C, but growth fell sharply above this temperature. The strains from the Bs lineage exhibited a lower growth rate at all temperatures but could still sustain growth 2 °C above the Bc strains. Despite experiencing the same fluctuations in temperature in the sediment of the small Churince lagoon, the two lineages could be easily discerned by their norm of reaction, suggesting differences in phenotypic plasticity.

**Figure 3.**
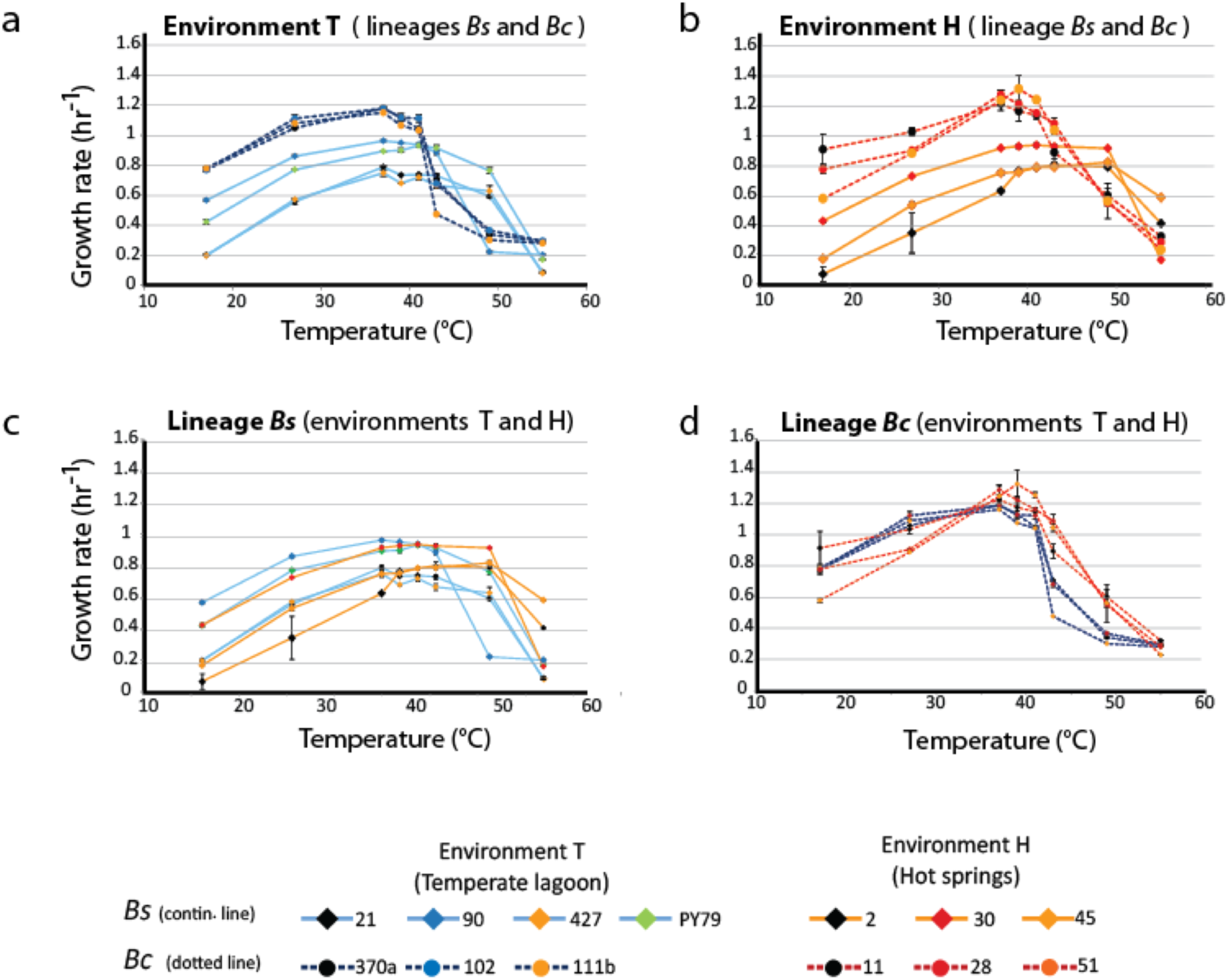
Lineage-specific phenotypic plasticity as a response to temperature. a. Norms of reaction to temperature of the B. cereus and B. subtilis lineages from the Temperate lagoon. b. Norms of reaction to temperature of the B. cereus and B. subtilis lineages from the Hot-springs. c and d. Combined data from curves in a and b, to highlight similarities in the response within the B. subtilis (c) and B. cereus lineage (d). Dashed lines, B. cereus lineage, continuous lines, B. subtilis lineage. Dark and light blue, Temperate lagoon strains. Red and orange, Hot-springs strains. Response was evaluated at temperatures 17, 27, 37, 43, 46, 49, 55 °C.

### Norms of reaction to temperature, a trait that differentiates the *Bacillus* lineages that co-occur in the hot-spring

Fig. 3b shows the profiles of the norms of reaction to temperature for the strains from the hot-spring. Norms of reaction to temperature of the strains belonging to the Bs lineage were more similar among them while those of the Bc lineage closely resembled each other. A higher selective pressure to temperature in a hot-spring did not lead to convergence of the two lineages in response to temperature; it seems thus that the distinct lineage-specific norm of reaction to temperature is a “stable” trait. As observed for these lineage strains from the temperature lagoon, a higher growth rate of the Bc lineage strains, followed by an abrupt drop was observed compared to that of the Bs strains, for which their growth pattern stretched smoothly towards higher temperatures.

### Evolution of tolerance to higher temperature of strains from the hot-spring

The strains of the Bs lineage in the H environment exhibited a wider range of temperature tolerance than those of the Bc lineage. Strains of Bs lineages sustained growth rate at higher temperature, and they reached a plateau and maintained the same growth response for a wide range of temperatures, to the point that no single optimal growth temperature could be defined. Growth only dropped at temperatures close to 50 °C (Fig. 3c).

In contrast, the strains from the Bc lineage from the hot-spring could reproduce more efficiently than those of the strains from the Bs lineage through all the temperature spectrum tested, until the temperature reached 42 °C, and then an abrupt drop in growth ensued. This can be clearly observed when comparing Fig. 3c, for lineage Bs, with Fig. 3d, for lineage Bc, as these graphs combine the norms of reaction from the T and H environment. Clearly, both Bs and Bc strains from the H environment exhibited higher tolerance for growth at temperatures above 42 °C, than those from environment T, suggesting that the strains have adapted to grow at a higher temperature. However, tolerance to temperature does not exceed more than a couple of °C more than the tolerance exhibited by the strains from the T environment. Even though the hot springs measured temperature fluctuate from 46 to 70 °C, these lineages do not exhibit plasticity to grow at temperatures above 45 °C. Noticeably, there is a tendency for strains from environment H to grow less at temperatures below 37 °C. A co-variance of higher temperature tolerance with a lower tolerance at lower temperature suggests a trade-off in phenotypic plasticity. The fact that each lineage exhibited a particular pattern and none of the strains in the Bc lineage could grow beyond 45 °C, supports the concept that the degree to which an organism can alter its phenotype is governed by its genetic architecture, that delimits the range of environmental conditions to which it can adapt.

### Thermal niche of strains from hot-springs and from the temperate lagoon

Fig. 4 is a summary of the measured parameters for both mesophilic and temperature-tolerant stains in both lineages, including thermal niche, optimum temperature and specific growth reached by individual strains at their optimal temperature (37 °C for most strains). The amplitude of the curve in the norm of reaction is the breath of the performance of the individual to a range of temperatures, while its thermal niche is the rank between the two lower and upper values of 75% of maximal growth at an optimal temperature, as described by (3). Regardless of lineage, all strains from the hot-spring exhibited a shift in their capability for growth at a higher temperature. For two strains in the Bc lineage from the H environment (strains Bc-H-51 and Bc-H-11), the extension in the capacity for growth at higher temperature seemed to impose a trade-off for growth at the lower temperature, and only strain Bc-H-28 exhibited increased tolerance without trade-off at low temperature. One of the strains from the Bs lineage from environment H (Bs-H-2) also exhibited a markedly lower capacity for growth at a lower temperature. All other strains from environment H exhibited a similar capacity for growth at a lower temperature as those from environment T, suggesting that they possessed the ability to grow at a wider range of temperatures.

**Figure 4.**
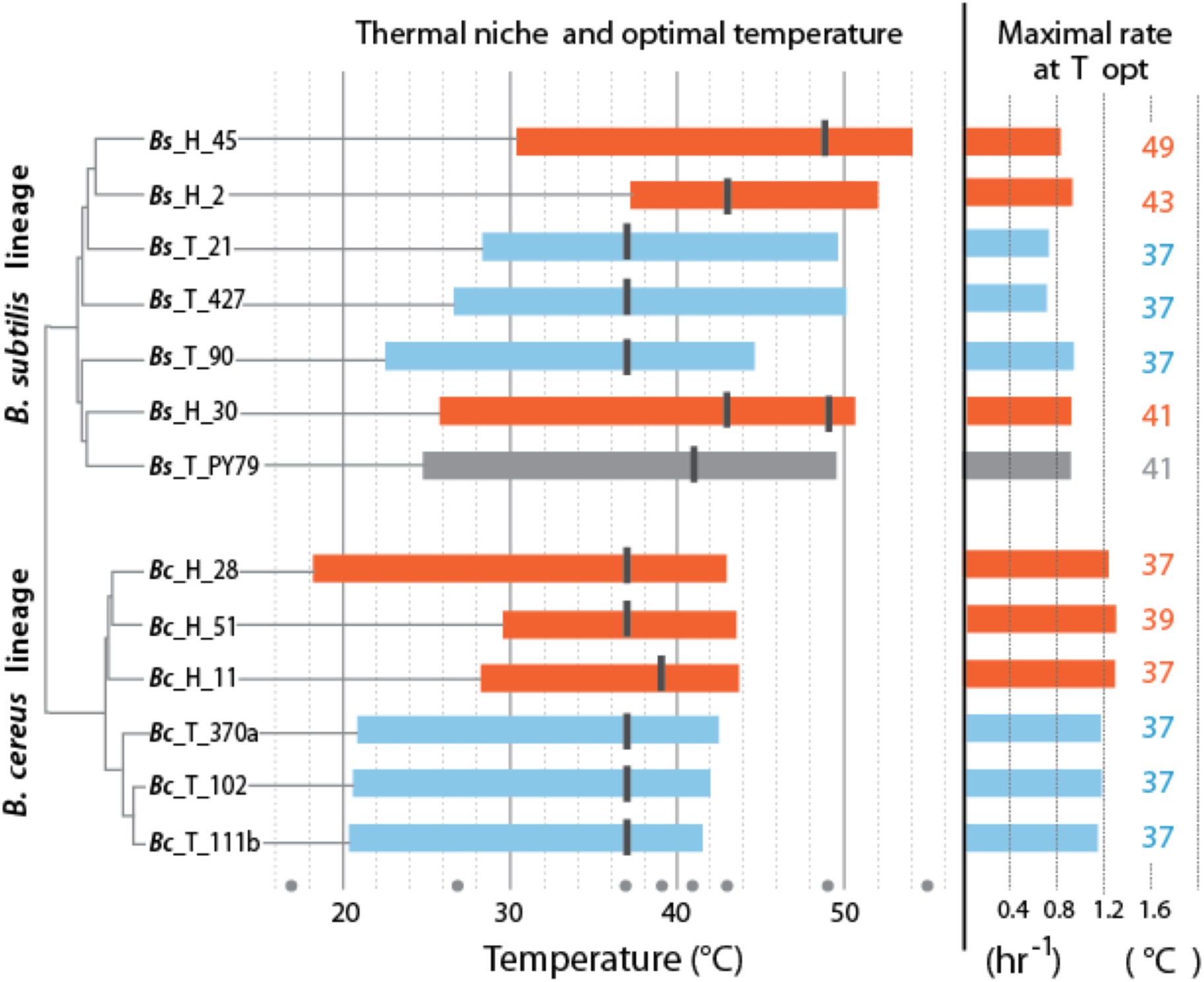
Thermal niche of the B. cereus and B. subtilis lineages from the temperate lagoon and the Hot-springs. Thermal niche is simplified as the range between the two x-values of 75% of maximal growth at optimal temperature as described by (3) and depicted in Fig. 1b. Rectangles depict range at or above 75% of maximum growth of strains from the Temperate lagoon (blue) and the Hot-springs (red). Black bars across the rectangle indicate the optimal temperature for growth (strain Bs H 30 exhibits maximum growth at two temperatures). Gray dots, temperatures of evaluation 17, 27, 37, 43, 46, 49, 55 °C. Maximum growth rate at optimal temperature for each strain is depicted to the right.

The optimal temperature for growth for all strains from both the Bc and Bs lineages that evolved in the T environment was 37 °C. Notably, all strains in the Bs clade from the environment H has a shifted optimal growth temperature to 43 °C (Bs-H-30 and Bs-H-2) and even to 49 °C (Bs-H-45). Interestingly, Bs-H-30 exhibited a plateau of optimal growth with a second optimal peak at 49 °C. It is intriguing that the laboratory strain, Bs PY79, exhibited a wide range of growth and even an optimal growth at 41 °C. The greater tolerance to temperature of B. subtilis compared to B. cereus lineage agrees with data obtained on semi-solid medium (Fig 2 and Supplementary Fig. 1).

It is evident that the strains from the Bc lineage from the H environment exhibited only 1 to 2 °C advantage in temperature tolerance and increased minimally their maximum capacity for growth. However, the specific growth rate of the Bc lineage strains, from either environment was always higher than that of strains from the Bs lineage, and this growth ability seems to be a trait of the species (Figure 4). This was also observed in the formation of larger colonies on plates (Fig. 2 and Supplementary Fig. 1).

Noticeable, within the Bc lineage, both optimal growth temperature and maximum optical density reached by the different strains measured at 37 °C, was similar to that of their counterparts from the temperate lagoon. The growth dynamics of Bc lineage isolates from the H environment do not exhibit the phenotypic plasticity expected for an organism from a constant environment above 46 °C. A graphic of the grouped data of the T and H strains from each of the lineages shows a clear lineage-specific norm of reaction to temperature (Fig. 5a), with the only point of convergence at 43 °C. The Bs lineage showed higher plasticity for growth at a higher temperature (strains from the T environment and, as expected, those from environment H). The strains within each lineage conserved a characteristic pattern in the norm of reaction that did not converge in their shared environments (Fig 5a). No statistical differences were observed when strains from T and from H were combined (Figure 5b) to compare environments, suggesting that species lineage exhibited a stronger signal in plasticity than the environment.

**Figure 5.**
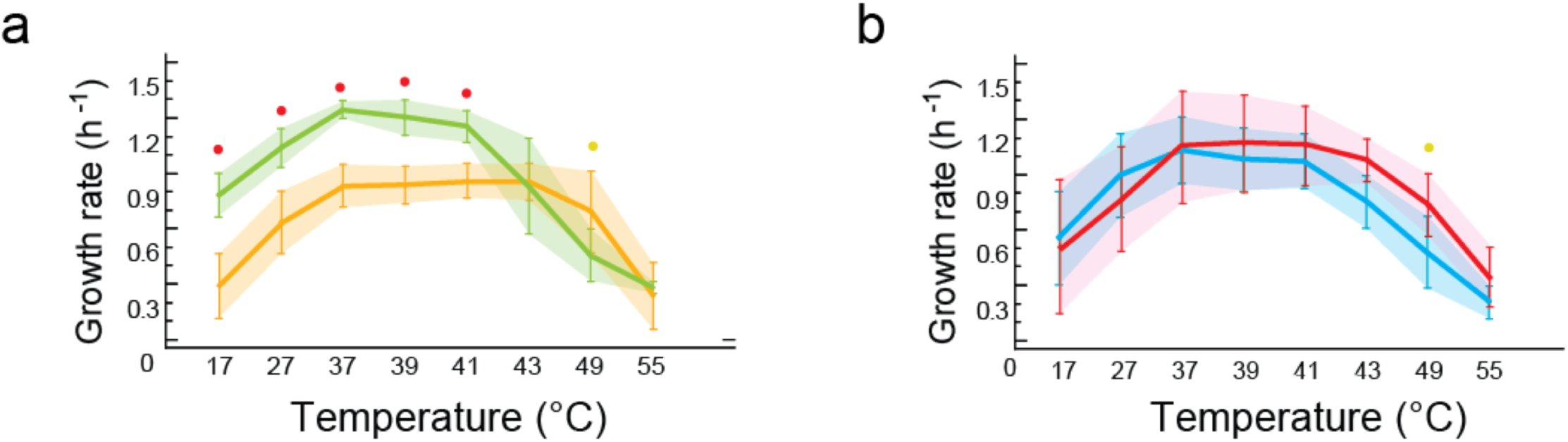
Phenotypic plasticity is constrained by genetic lineage a. Statistical analysis of growth rates of grouped-lineages. B. subtilis yellow, and B. cereus in green. Significant differences were obtained for growth response at five temperatures (red dots at 17, 27, 37, 39, 41°C). b. Statistical analysis of growth rates of grouped-by environment. T in blue, H in red. No significant differences in growth were observed when comparing data grouped-as-environments. Each curve represents the mean of growth rates for a set of strains. An ANOVA in R package 3.6.2 (at significance level at 0.05) was used to identify statistical differences in the double comparison, and intervals of confidence are shown with shaded area.

## DISCUSSION

It has been observed that realized niches for species in warm environments are closer to their physiological limits (5), but this has hardly been explored for bacteria. The presence of the same two *Bacillus* lineages in two contrasting environments, temperate and hot, provided the opportunity to evaluate the effect of evolutionary history on phenotypic plasticity as a response to temperature selection. A hot-spring constitutes selective pressure at what appears to be the edge of surviving temperature for mesophilic strains. Reaction norms, as a property of individual genotypes, allowed us to explore in a bacterial model the extent of phenotypic plasticity as the result of environmental history and the possible genetics constraints in two lineages.

There are reports of convergent evolution in bacteria when subject to experimental evolution (21). If bacteria isolated from the same environment responded in the same way to environmental challenges, despite differences in evolutionary history, this would suggest that prolonged evolution under stable conditions could lead to homogenous strategies to face environmental challenges. In our study, the evaluated species Bs and Bc lineages exhibited only intraspecific similarities rather than convergence patterns among the strains sharing a common selection regime (T or H). This suggests that deep evolutionary history of the individuals had set the genetic frame that determined their response to temperature, limiting their plasticity. This agrees with observations that plasticity to temperature can be regarded as a species trait (44).

Bs and Bc exhibited differences in their plasticity. The Bc lineage, with characteristic norms of reaction, showed an abrupt drop in growth after 42 °C. This behaviour has been called striking asymmetry and has been observed for reaction norms to temperature of many organisms, as performance increases and reaches an “optimal” level and then rapidly decreases near the lethal temperature (45). In contrast, the strains belonging to the Bs lineage exhibited a broader curve of tolerance and the strains from the H environment extended their tolerance to 47 °C, ten degrees above their optimal of 37 °C. The phenotypic plasticity of Bs seems to be superior in the isolates evaluated, including the laboratory strain PY79. It has been reported that Bs strain 168 can grow up to 52 °C (46). It is intriguing that the laboratory strain exhibits higher plasticity to temperatures it has probably not experimented, while Bc from the hot-spring did not become more tolerant to temperatures it has experimented possibly for centuries.

Another distinction between the lineages is the noticeable difference in growth and maximum growth rate (within their optimal range of temperature tolerance), with Bc strains exhibiting a faster growth rate than Bs. This also seems to be a lineage trait that did not change in either of the clades as long as it was evaluated within their thermal niche (Fig 4a). We had expected a decrease in the duplication time of the Bc strains, from the hot-spring as a possible trade-off of the ability to sustain growth at a higher temperature. This was not observed even at 44 °C. Its ecological strategy seems to be shifted towards faster growth, maybe to compensate that it can’t sustain growth at a higher temperature. This suggests that Bc is a specialist, with an r strategy, while Bs is a generalist, given the wide breadth of its thermal niche. These characteristics could have important implications when being part of a microbial community, particularly in constraining environments (47).

Our results suggest that sporulation is a form of plasticity that limits evolution. Since the Bacillus can sporulate, their thermal niche has to be defined for the vegetative and for the full developmental program leading to spore formation. For the *Bacillus* spp. (and other microorganisms), sporulation is the ultimate survival strategy allowing them to resist harsh environmental conditions (temperatures of 70 to 80°) for prolonged periods (48). However, sporulation is costly in time and energy investment and is a terminal differentiation decision. Endospore formation takes 8 h, and a genetic reprogramming that involves around 150 genes (49). Entering the sporulation process and remaining as spores would make the Bacillus numerically less competitive than if they could grow at a higher temperature in the hot-spring, where there may not be much of an environmental “intermission”. Although surviving is always a better option, this seems to be a case where sporulation limits genetic change and thus limits evolution towards heat tolerance in the vegetative stage. The spore has in fact been shown to evolve impressive features to shield DNA from damage (50). Although the *Bacillus* recovered from environment H could tolerate higher temperature in a vegetative stage than those from environment T, their optimal temperature for growth is still around 37oC and, surprisingly, they don’t grow above 50 oC. For the hot spring strains, phenotypic plasticity falls short at temperatures above 44 oC to 47 oC. With a limited thermal maximum for vegetative growth, these *Bacillus* probably survive in the hot springs as spores. If, as it has been suggested, under conditions in which plasticity is favoured, genetic variation can be limited (9), this might be a good example of this situation, as sporulation could limit the selection of tolerance in the vegetative phase. In addition to this possibility, could it be that no further tolerance to temperature can evolve in te vegetative stage, that the limit has been reached in these lineages? Our data showed that strains from environment H, as expected, were able to tolerate higher temperature for growth. This is consistent with data from experimental evolution studies using temperature as a selective environment and with data of bacterial isolates from natural environments. Experimental evolution work has been done mainly in *E. coli*. Populations evolved increased competitive fitness in the thermal regime that they experienced during the experiment (51), (22). The results from Bennett *et al*. (51) showed that *E. coli* strains evolved at 42 °C, can shift their tolerance towards higher temperature. On the other hand, experimental evolution of E. coli populations evolved for 20,000 generations at 37 °C were used to explore whether evolutionary adaptation to one particular environment leads to loss of performance in alternative environments. It was observed that improved performance at moderate temperature reduced V_max_ at extreme temperature (52).

Regarding bacteria from natural settings, Bronikowski *et al*. (3) did not observe variation in growth profiles for *Salmonella* or *E. coli* (comparisons within groups) isolated from turtle populations (undergoing natural changes in season temperatures) and from squirrels. Notwithstanding the lack of overlap between temperature ranges of different seasons, the breadth of all isolated strains were similar no matter what host they came from. On the other hand, the work of Sikorski et al. (53) showed increased tolerance to temperature among *Bacillus simplex* species isolated from a southern hill, that received more solar radiation and is consequently warmer and dryer, compared to strains from the northern hill. They also observed that the strains more tolerant (*B. simplex*) to temperature did not have a reduced capacity to grow at a lower temperature Sikorski and Nevo (53).

Can mesophilic *Bacillus* strains being exposed to strong selection at the limit of their temperature tolerance evolve thermophilic features or have they reached their temperature tolerance limit? Given their importance in food safety, several works have evaluated temperature tolerance in the *B. cereus sensu lato*. All in all, there are no examples of thermophilic strains in this group. Interestingly, seven major phylogenetic groups are now described that share a particular ‘ecotype’ structure in which each phylogenetic group exhibits its proper range of growth temperature and is for this reason associated with particular thermal niches. Clearly, only the *Bacillus cytotoxicus* (group VII) is thermotolerant (54). Interestingly, a trade-off is evident in strains that are more tolerant, as they exhibit less capacity to grow at a lower temperature (55). Group VII strains in this phylogenetic scheme are now recognized as a novel species ‘‘*B. cytotoxicus*” having a clearly distinguishing moderate thermotolerant phenotype (56) and being genetically distant from more than 200 *B. cereus* examined (57). Interestingly, even spore inactivation temperatures are different among the B. cereus groups, with those that tolerate more temperature in the vegetative stage also requiring more temperature for spore inactivation (58), (55). Several works have evaluated the temperature for spore inactivation in different *Bacillus* species. Den Besten et al. reviewed data for *Bacillus* spp. collected from different places, mainly food sources. Although there is variation in heat inactivation of spores, those of Bs lineage seem to be more tolerant than those of Bc lineage (59).

It has been suggested that in ectotherms in a rapidly changing environment there is a trade-off between maximal performance, particularly in thermal specialists in contrast to thermal generalists (45). Miller and Castenhotz (60) evaluated *Synecococcus* from hot-springs and noted that as the upper-temperature limit for growth was extended, an even larger shift upward in the minimum temperature was observed, leading them to suggest that increases in thermal specialization resulted in a decrease in the overall temperature range for growth. In this work, only for some strain we observed that higher performance at temperatures beyond 37 °C came at a cost to growth at a lower temperature.

Data from different reports is consistent with our observation that genetic architecture is a defining element to temperature tolerance. Lineage can be easily identified through its norm of reaction in both environments, the genetics behind the response to temperature may constrain changes in phenotypic plasticity. Environmentally induced plastic phenotypes are thought to be controlled by gene regulatory networks (61) that often have a common regulation. Environment-specific gene expression has long been appreciated to underlie plasticity in prokaryotes (see for example (62)). The heat shock response was the first regulatory system discovered and is considered one of the fundamental systems concerning general stress (63). Some effectors of the heat shock proteins are highly conserved in all three domains of life. Bacteria and lower eukaryotes share conserved families of chaperones and maybe also conserve the complexity of thermal systems such as chaperone networks (37), which as a system may be slower to evolve. This may be the reason for the resistance to change in genetic lineages and would explain why there is a correlation in plasticity to temperature as a function of the organisms’ genetics, and not of the environment. It is also possible that environmental parameters other than temperature (e.g., nutrient availability, pH, and interactions with other microorganisms) may influence the overall fitness of these organisms and thus limit their plasticity.

In summary, phenotypic plasticity of temperature tolerance (thermal acclimation) is considered an important component of the evolutionary response to variable temperatures and specifically as a relevant response to climate change (4). Understanding how organisms respond and adapt to novel environments is critical to our efforts to conserve biodiversity and maintain ecosystem function. It was not expected that the heat tolerance phenotype of the Bacillus in the hot spring would not match its habitat’s temperature. Phenotypic plasticity seems to be a lineage trait, each of the *Bacillus* lineages seems to possess a distinctive reaction norm to temperature and possibly a rigid genetic architecture that limits convergence. It is possible that substantial molecular changes may be required to increase upper thermal limits for the *Bacillus* and that the observed limited tolerance to temperature reflects evolutionary constraints. If this scenario is also true for other bacteria, the limited potential to change their thermal limits should be a strong warning particularly within the context of an average predicted temperature increase of 2–4 °C for mid-latitude populations over the next few decades. It would be interesting to test the hypothesis of the genetic constraints on thermal tolerance by subjecting the *Bacillus* strains from the temperate environment to experimental evolution to find their thermal boundaries.

## Conclusion

Despite sharing the same environment (hot or temperate) the evaluated *Bacillus* strains from the two lineages do not converge in their norms of reaction to temperature. Deep evolutionary differences define the genetic possibilities of plasticity to temperature, such that the norms of reaction to temperature can be considered a strong lineage signature for the Bc and Bs strains analyzed. Sporulation allows the hot-spring *Bacillus* strains to exceed what would be their temperature tolerance limit in the vegetative stage, and although the spore state allows their survival, it may reduce opportunities to evolve higher tolerance. However, given that the thermal niche was observed to be shifted only a few degrees toward more tolerance to temperature, it is possible that there is a genetic architecture constraint and that these lineages have reached their tolerance limit. The reduced plasticity exhibited by these bacterial lineages should be a warning for the limited capability, even of bacteria, to adjust to climate change.

## Acknowledgments

Fund for this work came from Consejo Nacional de Ciencia y Tecnología (Conacyt) Básica 2014 CLAVE 220536 to G. O.A. E.H. acknowledges fellowship from Consejo Nacional de Ciencia y tecnología (Conacyt). We acknowledge the technical help of many under-graduate students. We are grateful to Dr. Gabriel Moreno for many fruitful discussions.

