## Supplementary material for "Phenotypic plasticity and evolution of thermal tolerance in two lineages of bacteria from temperate and hot environments": Supplemnary Figures 1 and 2

**Supplementary figure 1.** Colonies of *Bacillus* strains grown on Marine Medium at different temperatures. Colonies of selected strains were streaked out on semisolid Marine medium and incubated for 24 to 48 h at 37, 44, 50 and 55 °C. The plates were photographed after a 24 h incubation and returned to the incubator at the same temperature for another 24 h incubation.

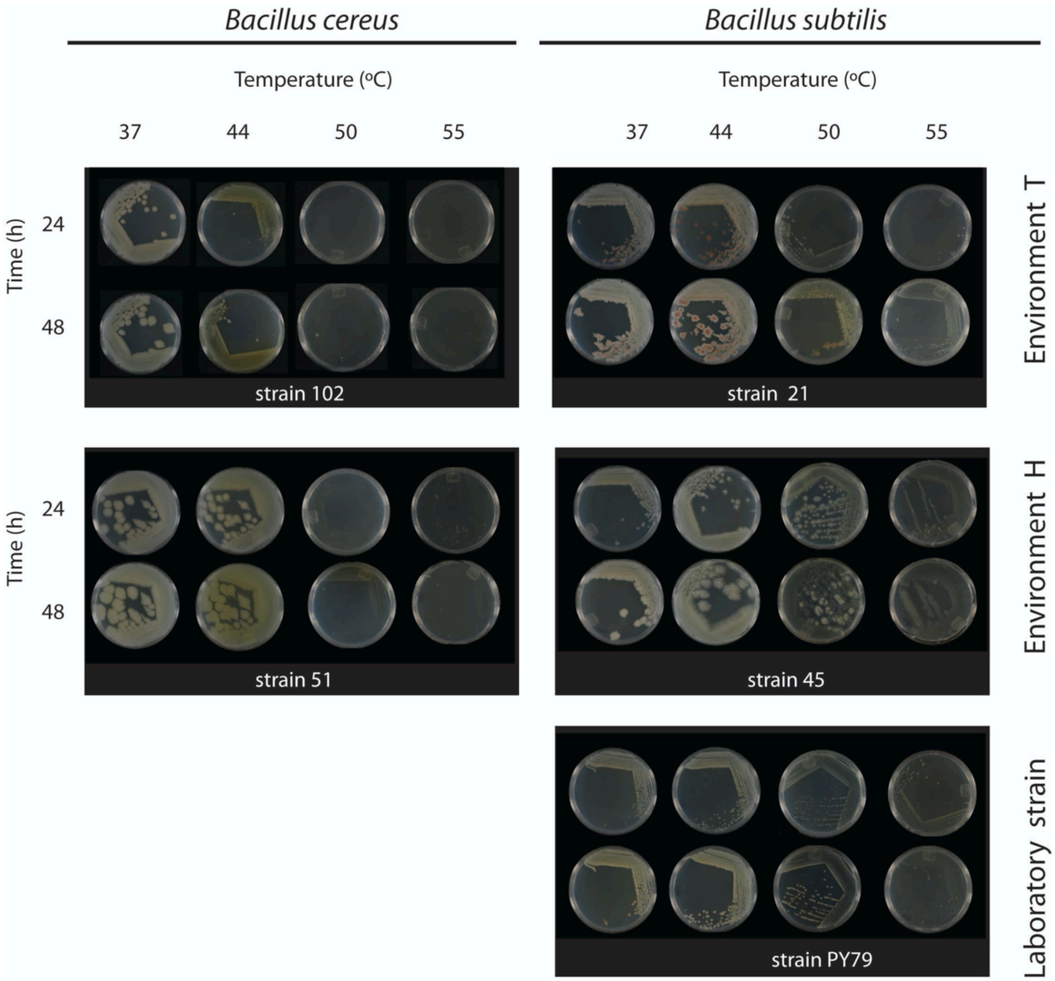

**Supplementary figure 2.** Phylogenetic analysis based on the 16S rRNA gene of *Bacillus* strains from our collection from the Hot-springs and the temperate lagoon of Churince. A maximum likelihood phylogenetic analysis was done based on the 16S RNA gene to select strains that belonged to the either the *Bacillus cereus sensu lato* (lineage B) or the *Bacillus subtilis sensu lato* (lineage A).

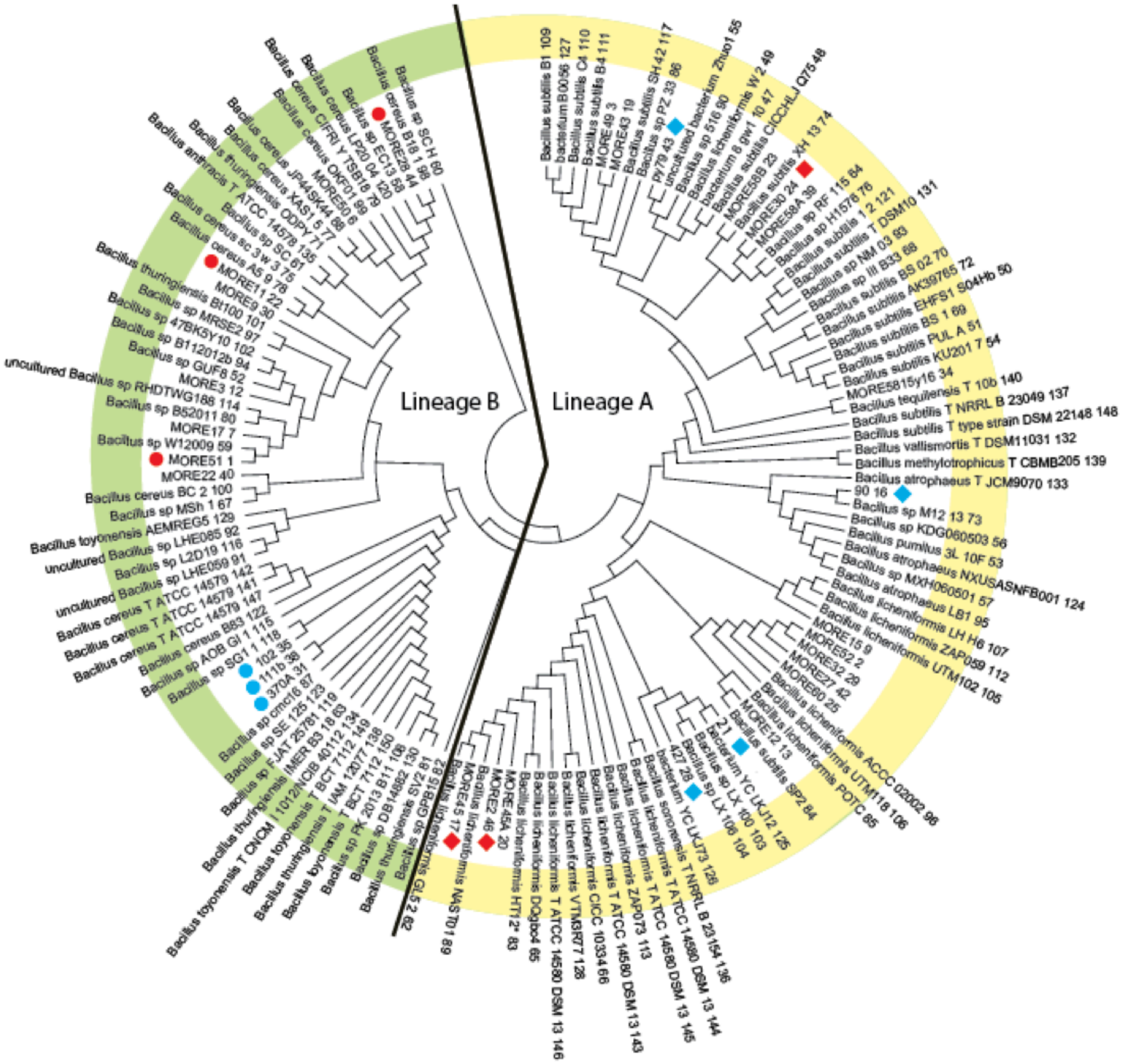
